# The Arabidopsis leucine-rich repeat receptor kinase MIK2 is a crucial component of pattern-triggered immunity responses to *Fusarium* fungi

**DOI:** 10.1101/720037

**Authors:** A.D. Coleman, L. Raasch, J. Maroschek, S. Ranf, R. Hückelhoven

## Abstract

*Fusarium* is a genus of fungi causing severe economic damage in many crop species exemplified by *Fusarium* Head Blight of wheat or Panama Disease of banana. Plants sense immunogenic patterns (termed elicitors) at the cell surface contributing to disease resistance *via* the activation of pattern-triggered immunity (PTI). Knowledge of such elicitors or corresponding plant immunity components is largely lacking for *Fusarium* species. We describe a new peptide elicitor fraction present in several *Fusarium spp.* which elicits canonical PTI responses in *Arabidopsis thaliana* but depends on a currently unknown perception mechanism. We therefore employed a forward-genetics screen using Arabidopsis plants containing a cytosolic calcium reporter (apoaequorin) to isolate *fere* (*Fusarium Elicitor Reduced Elicitation*) mutants. The *fere1* mutant showed impaired PTI marker responses to an enriched elicitor fraction derived from *Fusarium oxysporum* but normal responses to other fungal elicitors. We mapped the causal mutation to the receptor-like kinase MIK2 (MALE DISCOVERER1-INTERACTING RECEPTOR LIKE KINASE 2) with a hitherto undescribed role in PTI pathways but documented functions in other cell surface signalling pathways. The strength of the phenotype in *fere1* and independent *mik2* mutants supports that MIK2 is a new key component in sensing *Fusarium*. *Fusarium* elicitor responses also partially depend on PTI signalling components known for other cell surface elicitor responses such as BAK1, BIK1, PBL1, FERONIA, LLG1 and RBOHD. This shows that Arabidopsis senses *Fusarium* by a novel receptor complex at the cell surface that feeds into common PTI pathways and positions MIK2 as a central player that potentially integrates plant endogenous signals with biotic and abiotic stress responses.

## Introduction

Plants have evolved diverse pattern-recognition receptors (PRRs) at the cell surface to detect microbial invaders and form multipartite PRR complexes to initiate and regulate intracellular signalling pathways. Molecular patterns from diverse origins (self or non-self; collectively termed elicitors) can be perceived by plant PRRs contributing to quantitative disease-resistance via the activation of pattern-triggered immunity (PTI). *Fusarium* is a large genus of fungi causing severe economic damage in many cultivated plant species and although several studies have demonstrated mechanisms to detect and respond to *Fusarium*, knowledge of elicitors from *Fusarium* or subsequently induced plant immune-signalling is sparse. To improve mechanistic understanding of *Fusarium*-triggered PTI, we employed a forward-genetics screen in Arabidopsis to isolate a mutant strongly impaired in PTI responses to an elicitor fraction from *Fusarium* spp. but fully responding to other fungal elicitors. We identified a causal mutation in the leucine-rich repeat receptor kinase MIK2 (MDIS1-INTERACTING RECEPTOR LIKE KINASE2), which was previously described in the context of abiotic and biotic stress resistance, sexual reproduction and cell wall integrity sensing. Importantly, MIK2 was also shown to contribute to Arabidopsis resistance to *Fusarium oxysporum*. Here, we now describe a novel and crucial function of MIK2 in pattern-triggered immunity.

## Results and Discussion

Much of what we currently understand about PTI in plants has been established in *Arabidopsis thaliana* where several PRRs or associated signalling components have been described. To determine the feasibility of identifying *Fusarium*-relevant PTI components by forward genetics, we characterised the hallmark PTI response of cytosolic elevation of calcium concentration in Arabidopsis to crude *Fusarium graminearum* elicitor (FGE) or *Fusarium oxysporum* elicitor (FOE) fractions derived from fungal mycelium by the use of apoaequorin reporter plants (Knight *et al.* 1991; Ranf *et al.* 2011) (Figure 1a). Similarly prepared fractions from additional *Fusarium* species including *F. culmorum*, *F. avenaceum* and *F. langsethiae* also gave comparable responses in Arabidopsis, indicating the presence of similar molecular patterns. Chitin is a potent fungal elicitor in plants. The chitin elicitor receptor-like kinase 1 mutant (*cerk1*) lacks an essential component of chitin-mediated PTI in Arabidopsis (Miya *et al.* 2007) yet showed a wildtype response to crude *Fusarium* elicitor preparations such as FGE, indicating that chitin is not a major contributor of the responses we observe (Figure 1b). As crude mycelial fractions likely contain multiple elicitors and thereby initiate variable PTI responses, we enriched an elicitor fraction from *F. oxysporum* by binding and elution from ion exchange resins and termed the fraction EnFOE (enriched *F. oxysporum* elicitor). EnFOE gave robust and consistent responses with multiple PTI marker assays in Arabidopsis, including cytosolic Ca^2+^ elevations, reactive oxygen species (ROS) production, mitogen-activated protein kinase (MAPK)-phosphorylation and defence-marker gene activation (Figure 1c-f).

**Figure 1:**
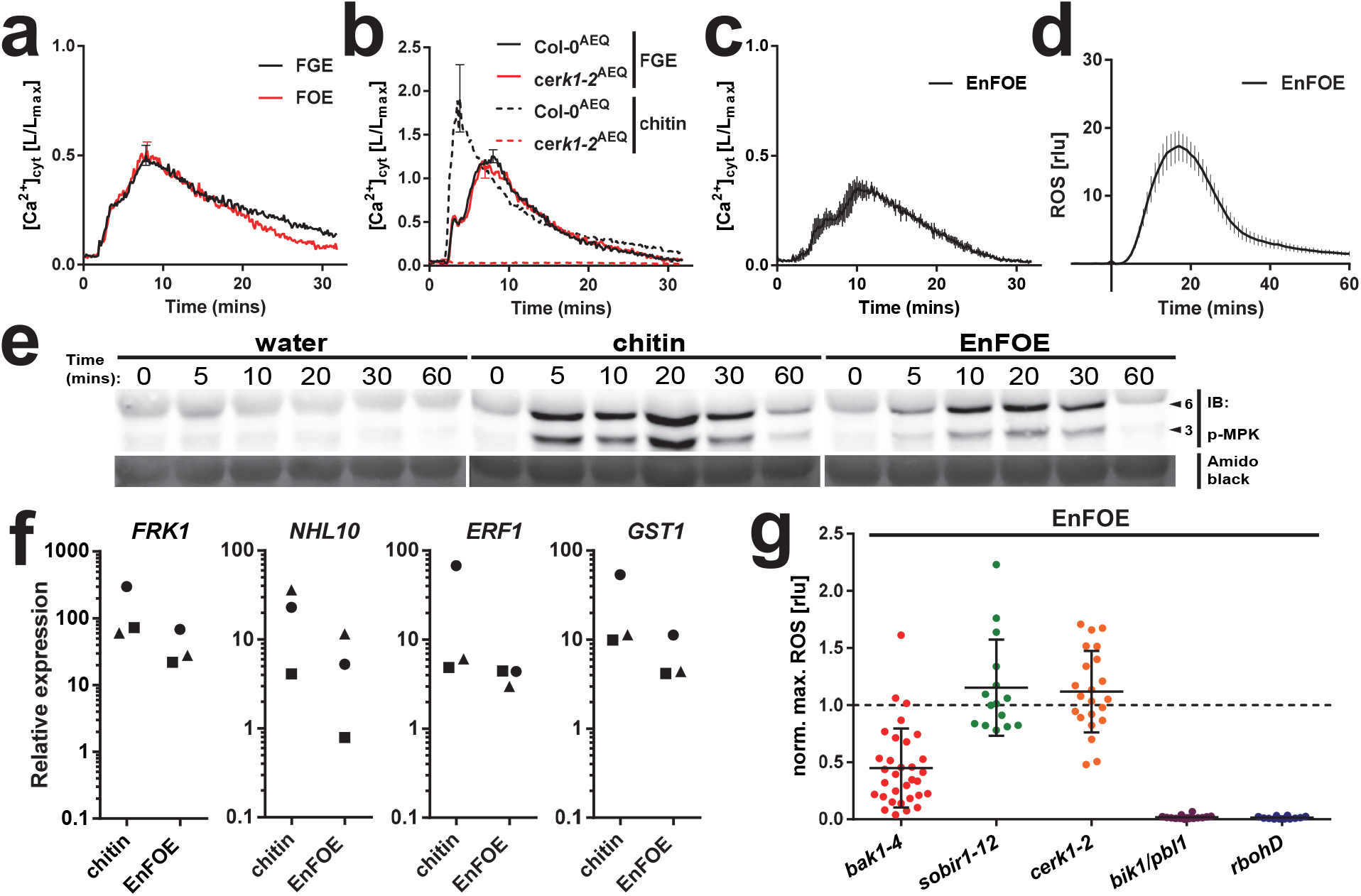
An elicitor enriched from *Fusarium* species elicits hallmark PTI responses in Arabidopsis partially dependent on known signalling components. [**a**] Cytosolic Calcium quantification ([Ca^2+^]_cyt_) kinetics in Col-0^AEQ^ seedlings after elicitation with crude *F. graminearum* elicitor or *F. oxysporum* elicitor fractions (FGE/FOE; derived from 1 mg mycelium/mL). [**b**] [Ca^2+^]_cyt_ kinetics in Col-0^AEQ^ or *cerk1-2*^AEQ^ seedlings after elicitation with FGE (2.5 mg mycelium/mL; full line) or chitin (60 µg/mL; dotted line). [**c**] [Ca^2+^]_cyt_ kinetics in Col-0^AEQ^ after elicitation with the enriched *F. oxysporum* elicitor fraction (EnFOE; 0.5 µg/mL). [Ca^2+^]_cyt_ data represent mean values measured in individual seedlings over 30 minutes after elicitation with error bars indicating s.d. at the maximum of the curve (a-b) or at each measurement point (c) (*n*=4). [**d**] ROS accumulation kinetics in Col-0^AEQ^ leaf discs after elicitation with EnFOE (0.5 µg/mL). Data represent mean relative light units [rlu] measured from individual leaf discs with error bars representing s.d. at each measurement point (*n*=4). [**e**] Immunoblot analysis (IB) of phosphorylated MPK6 (6) and MPK3 (3) (arrowheads at right margin; p-MPK; upper panel) in Col-0^AEQ^ seedlings after treatment with water, chitin (75 µg/mL) or EnFOE (0.5 µg/mL) over the time series stated, including Amido black staining of total proteins (loading control; lower panel). [**f**] Expression of PTI-marker genes in Col-0^AEQ^ seedlings two hours after treatment with chitin (75 µg/mL) or EnFOE (0.5 µg/mL) relative to expression in water-treated plants (set at 1; symbols represent three independent biological replicates; target genes normalized to expression of *Ubiquitin*). [**g**] Maximum ROS accumulation measured in individual leaf discs (each plotted point) of various PTI-relevant mutant genotypes over 60 minutes after elicitation with EnFOE (0.5 µg/mL) normalized to the mean maxima of Col-0^AEQ^ plants (set at 1; dotted line) with error bars representing s.d. (*n*=12–32). Data shown is representative from at least three independent experiments with similar results (a-e) or normalized pooled data from at least three independent biological replicates (g).

Depending on receptor class and ectodomain structure, PRRs require various additional signalling partners for full signalling capability. To determine whether known PTI components are necessary for recognition of EnFOE, we tested EnFOE-induced PTI responses in a range of mutants affected in known PTI signalling components. The receptor-like kinase SERK3/BAK1 is a central regulator of innate immunity in plants to immunogenic peptides and associates with multiple PRRs to initiate signal transduction in a partially redundant manner with other SERK family proteins (Liebrand, van den Burg, and Joosten 2014). EnFOE-induced ROS production was approximately 50% reduced (Figure 1g) and noticeably delayed in *bak1-4* compared to WT plants (Figure S1). No EnFOE phenotype was observed in additional SERK mutant lines (Figure S1), suggesting that BAK1 is involved in EnFOE-triggered immune signalling. However, PTI responses to EnFOE were shown to be independent of SOBIR1 (a core component of receptor-like protein type receptors) as well as CERK1, as shown previously for crude *Fusarium* elicitor fractions (Figure 1g). Nevertheless, BIK1 and PBL1, cytoplasmic kinases which are involved in many PTI pathways, and the ROS producing NADPH oxidase RBOHD are required for full ROS responses to EnFOE (Figure 1g). Hence, there is convergence of responses to EnFOE with those to other described elicitors. To further test for recognition of EnFOE by known PRRs, we tested EnFOE responses in additional mutant lines and confirmed that recognition is independent of RLP23, RLP30 and RLP42, each involved in detection of fungal elicitors (Albert *et al.* 2015; Zhang *et al.* 2013; Zhang *et al.* 2014), as well as RFO1, 2 or 3 (Figure S1), known to be involved in quantitative resistance to *F. oxysporum* in Arabidopsis (de Sain and Rep 2015). EnFOE-induced PTI responses could be abolished using a kinase inhibitor and first biochemical characterisation of EnFOE determined that activity is mostly proteinaceous with a typical dilution response profile (Figure S2). As determined through the elicitor preparation method, EnFOE is also heat-stable and likely above 14 kD in size (see methods). Although both roots and above ground tissues are responsive, EnFOE-triggered cytosolic Ca^2+^ elevations are higher in the root. This was similar to the chitin response and in contrast to *Fo*NLP response, which was hardly detectable in roots (Figure S3). Weak responses of the root were also reported for bacterial peptide elicitors elf18 and flg22, which are predominantly detected in above ground tissues (Ranf *et al.* 2011). Calcium elevations to nlp20 are exclusively detected in the meristematic zone of the root (Wan *et al.* 2018) which could explain the low responsiveness to *Fo*NLP we observe in root tissue.

As the precise molecular identity of the elicitor was not necessary for forward genetic approaches, we screened Arabidopsis (Col-0^AEQ^) EMS populations (similar to: Ranf et al. (2012)) for attenuated cytosolic Ca^2+^ response to FGE. We focused on mutants with strongly reduced responses to FGE but largely unaltered responses to chitin and flg22. This identified a mutant termed *fere1* (*Fusarium elicitor reduced elicitation 1*), which has partially reduced responses to crude FGE/FOE fractions, genetically supporting the presence of similar elicitors in different *Fusarium* species (Figure S4). Interestingly, EnFOE-induced PTI responses are abolished in *fere1* showing that we successfully enriched the immunogenic fraction of FOE (Figure 2a-b). We further confirmed that *fere1* is able to respond to the fungal elicitors chitin and *Fo*NLP, suggesting specificity to EnFOE-induced PTI. Interestingly, *fere1* shows slightly increased responses to these other fungal elicitors (Figure 2c).

**Figure 2:**
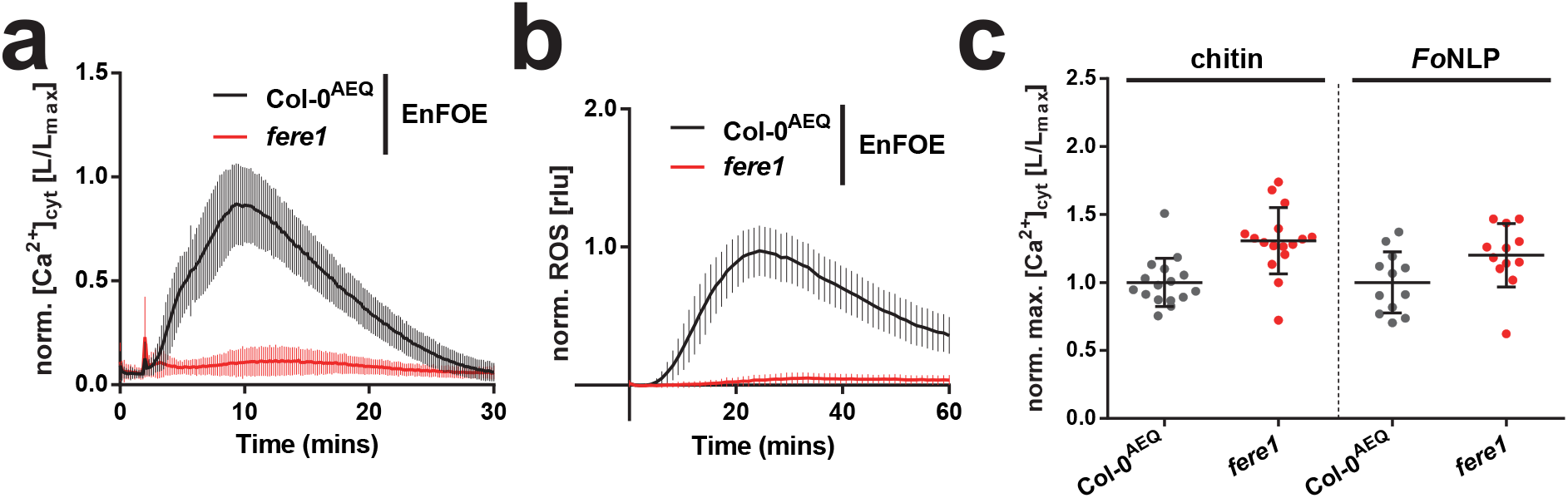
The *Fusarium elicitor reduced elicitation* (*fere1*) mutant is strongly impaired in PTI responses to the *Fusarium* elicitor. [**a**] Cytosolic Calcium quantification ([Ca^2+^]_cyt_) kinetics in Col-0^AEQ^ or *fere1* seedlings after elicitation with the enriched *F. oxysporum* elicitor fraction (EnFOE; 0.5 µg/mL). Data represent mean values measured in individual seedlings normalized to respective values from Col-0^AEQ^ plants with error bars representing s.d. at each time point (*n*=84). [**b**] ROS accumulation kinetics in Col-0^AEQ^ or *fere1* leaf discs after elicitation with EnFOE (0.5 µg/mL). Data represent mean relative light units [rlu] measured from individual leaf discs normalized to respective values from Col-0^AEQ^ plants with error bars representing s.d. at each time point (*n*=19–20). [**c**] Maximum [Ca^2+^]_cyt_ measured in individual Col-0^AEQ^ or *fere1* seedlings (each plotted point) over 30 minutes after elicitation with chitin (75 µg/mL) or *Fo*NLP (1 µM) normalized to the mean maxima of Col-0^AEQ^ plants (set at 1) with error bars representing s.d. (*n*=12–16). Data shown (a-c) is pooled from at least three independent biological replicates.

To identify the causal single nucleotide polymorphism (SNP) in *fere1*, we generated a *fere1* outcross population with ecotype Ler-0 and confirmed phenotypic segregation of a single recessive locus, which we subsequently mapped using insertion/deletion (InDel) markers (Salathia *et al.* 2007) to a region of approximately 0.4 MB on the short arm of chromosome 4 containing 121 gene loci. The mapping interval contained a gene for the LRR-RLK MIK2 (MALE DISCOVERER1 (MDIS1)-INTERACTING RECEPTOR LIKE KINASE2; also known as LRR-KISS). Because *MIK2* was the only gene in the mapping interval that encodes an obvious candidate PRR complex component by protein domain architecture, we sequenced *MIK2* in *fere1* and identified a SNP resulting in a premature W876STOP codon within the cytoplasmic kinase domain. A gene model representing the *fere1* SNP and various *mik2* alleles used in this study is shown in Figure 3a. MIK2 is a class XII LRR-RLK with described roles in cell wall integrity sensing, root growth, and response to abiotic and biotic stress (Julkowska *et al.* 2016; Van der Does *et al.* 2017). It is reported to heterodimerize with MDIS1 and MDIS1-INTERACTING RECEPTOR LIKE KINASE1 to mediate male perception of female defensin-like LURE peptides during pollen tube attraction (Wang *et al.* 2016). *MIK2* has one intron and the pre-mRNA is spliced into two forms (both of which are affected by the *fere1* SNP), with *MIK2.1* being up to 50 fold more abundant than *MIK2.2* (Van der Does *et al.* 2017). Three independent *mik2* mutant alleles show largely abolished PTI responses to EnFOE but not to chitin (Figure 3b). Furthermore, *mik2-3* is allelic to *fere1* which we tested by reciprocal crossing. In addition, we generated *fere1*-independent *mik2-3* mutants carrying the apoaequorin reporter (*mik2-3*^AEQ^) which showed a similar loss-of-function phenotype in response to EnFOE (Figure 3c). Furthermore, the PTI loss-of-function phenotype in *fere1* to EnFOE was fully complemented by ectopic *MIK2* expression under the control of its own promoter as confirmed by ROS and cytosolic Calcium response measurements (Figure 3d-e). This was similarly achieved by using the predominant *MIK2.1* splice form amplified from cDNA, as well as the full-length genomic region of *MIK2* with a C-terminal cMyc peptide fusion.

**Figure 3:**
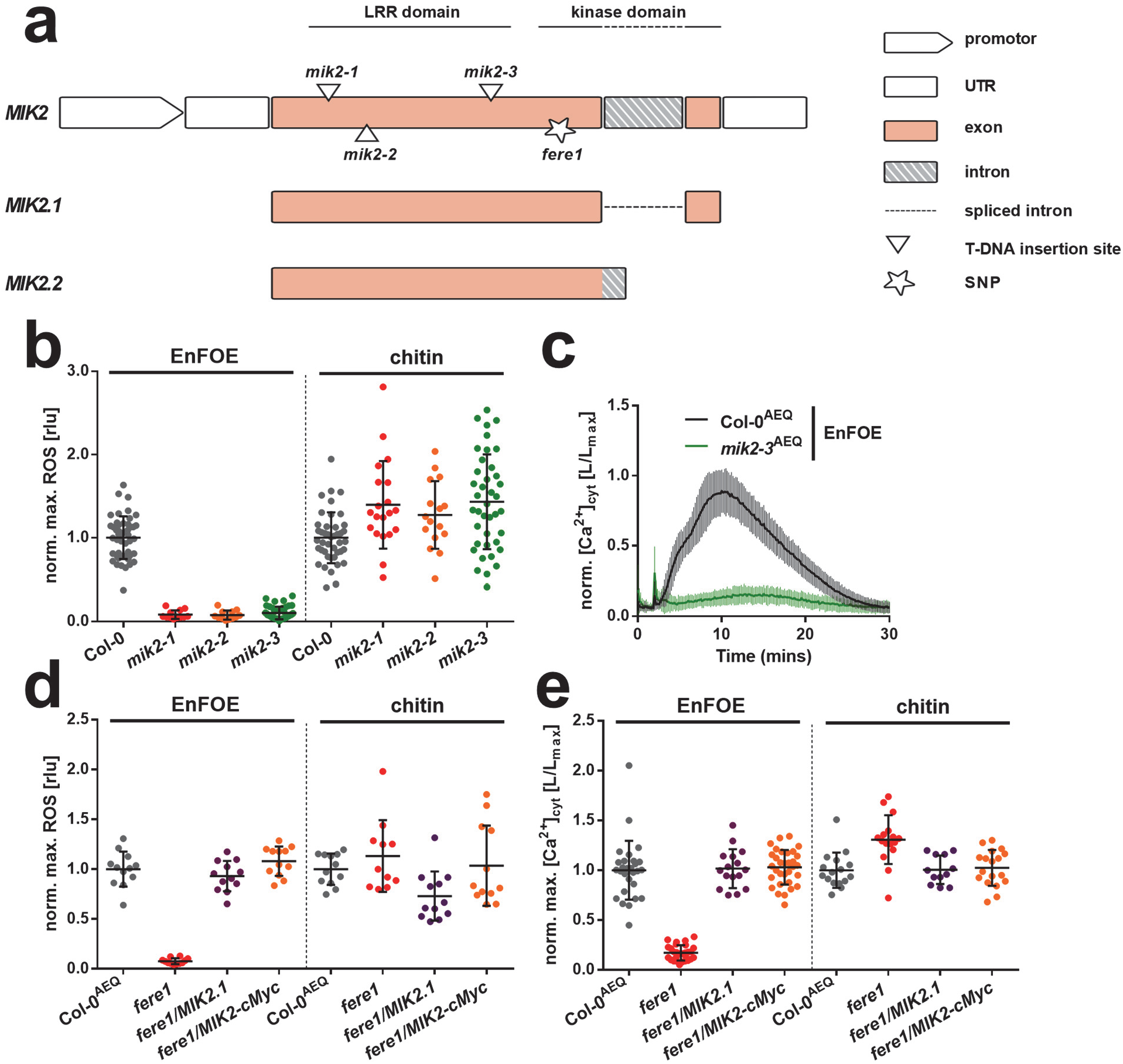
The *fere1* mutant contains a causal mutation in *MIK2*. [**a**] *MIK2* gene model showing predicted splice forms (*MIK2.1*, *MIK2.2*) and locations of *fere1* SNP and T-DNA insertions. **[b]** Maximum ROS accumulation measured in individual leaf discs (each plotted point) from Col-0 or three independent *mik2* T-DNA alleles over 60 minutes after elicitation with the enriched *F. oxysporum* elicitor fraction (EnFOE; 0.5 µg/mL) or chitin (75 µg/mL) normalized to the mean maxima of Col-0 plants (set at 1) with error bars representing s.d. (*n*=12–47). [**c**] Cytosolic Calcium quantification ([Ca^2+^]_cyt_) kinetics in Col-0^AEQ^ or *mik2-3*^AEQ^ seedlings after elicitation with EnFOE (0.5 µg/mL). Data represent mean values measured in individual seedlings normalized to respective values from Col-0^AEQ^ plants with error bars representing s.d. at each time point (*n*=40). [**d**] Maximum ROS accumulation measured in individual leaf discs (each plotted point) from Col-0^AEQ^, *fere1*, *fere1/MIK2.1* or *fere1/MIK2-cMyc* over 60 minutes after elicitation with the enriched *F. oxysporum* elicitor fraction (EnFOE; 0.5 µg/mL) or chitin (75 µg/mL) normalized to the mean maxima of Col-0^AEQ^ plants (set at 1) with error bars representing s.d. (*n*=11–12). [**e**] Maximum [Ca^2+^]_cyt_ measured in individual Col-0^AEQ^, *fere1, fere1/MIK2.1* or *fere1/MIK2-cMyc* seedlings (each plotted point) over 30 minutes after elicitation with EnFOE (0.5 µg/mL) normalized to the mean maxima of Col-0^AEQ^ plants (set at 1) with error bars representing s.d. (*n*=12–32). Data shown (b-e) is pooled from at least three independent biological replicates.

So far, MIK2 has not been implicated in PTI responses, but its expression has been shown to be regulated by pathogen infection or elicitor treatment (based on the Arabidopsis eFP Browser by Winter *et al.* (2007); Expression Sets 1007966202 and 1008080727). Notably, MIK2 is important for resistance against *F. oxysporum* which was previously linked to its role in cell wall integrity (CWI) monitoring (Van der Does *et al.* 2017). Our data now suggest that the additional role of MIK2 in detecting EnFOE to trigger PTI may be a major contributor towards resistance against *Fusarium* species and could therefore explain the higher susceptibility previously reported for *mik2*. PTI and CWI sensing are intimately interconnected (Hamann 2012; Wolf 2017; Engelsdorf *et al.* 2018) and it has for example been reported that *mik2* shows reduced isoxaben-induced cell wall damage (Van der Does *et al.* 2017; Engelsdorf *et al.* 2018). Thus, to determine whether CWI sensing generally interferes with sensing EnFOE, we tested EnFOE responsiveness in additional cell-surface receptor mutants with known roles in CWI e.g. HERKULES, WAK2, THESEUS and FERONIA (Engelsdorf *et al.* 2018). A WT-like response was observed in *herk1*, *wak2*, and two independent *the1* alleles (*-1* [null mutant] and *-4* [reported to be hypermorphic; Merz *et al.* (2017)]) to both EnFOE and chitin (Figure S5) showing that impairment in CWI sensing does not generally interfere with the response to EnFOE. Interestingly, a markedly reduced response was observed in *fer-4* and *llg1* mutants (a chaperone and co-receptor of FERONIA involved in perception of RALF peptides; Li *et al.* (2015); Xiao *et al.* (2019)) (Figure S5), indicating a role of FERONIA in EnFOE-induced PTI signalling consistent with its role in controlling various immune signalling processes (Stegmann *et al.* 2017). At this point, it remains open whether MIK2 is a PRR for peptides in EnFOE. However, the almost complete loss-of-function phenotype in diverse *mik2* alleles when elicited with EnFOE argues for a function as a PRR, co-receptor or essential regulator of elicitor perception. Although the strong loss-of-function phenotype of *fere1* and other *mik2* alleles in response to EnFOE argues against redundancy of this response, we also tested mutants in *MIK2-LIKE* (60% amino acid identity to MIK2) (Van der Does *et al.* 2017). The *mik2-like* mutant plants did not show a reduced ROS response to EnFOE however (Figure S1), suggesting that MIK2-like does not fulfil a similar function as MIK2 in PTI.

The nature of immunogenic peptides in EnFOE remains unknown for now. However, the role of MIK2 in sexual compatibility sensing via LURE peptides suggests that MIK2 can perceive endogenous and exogenous signals. With its additional function in CWI sensing and salt stress resistance, MIK2 could act as an integrator of diverse signals for coordinated plant response to the environment. This does not exclude that MIK2 could also function as a direct peptide receptor. Comparably, the RLK FERONIA, which has multiple functions in plants, was shown to be also a receptor for RALF peptides (Stegmann *et al.* 2017; Xiao *et al.* 2019). As MAMPs can be highly conserved patterns that a microbe cannot easily dispense of, it will be important to determine how widespread EnFOE-like activity is within the fungal kingdom. It has been recently underlined that identification of genetic components with key functions in response to complex elicitors is critical towards the identification of the smallest active compounds in such elicitors. For instance, the LORE receptor was crucial for further chemical dissection of the plant PTI response to enriched lipopolysaccharide fractions from *Pseudomonas* species and guided the identification of medium chain length 3-hydroxy fatty acids as the actual bacterial elicitors (Kutschera *et al.* 2019). We therefore suggest that the identification of MIK2 as a key component in PTI responses to EnFOE will foster future work on identification of immunogenic peptides from *Fusarium* species and on identification of further PTI components specific for EnFOE responses. The latter could be identified via physical and/or genetic interaction with MIK2 and may disclose the nature of the PRR complex responsible for PTI responses to EnFOE. Findings from the model Arabidopsis will speed up the process of identifying similar components in *Fusarium*-relevant crop plants such as banana, wheat, barley, maize or soybean, where comparable genetic resources are not fully available and informative PTI assays are less well established. Finally, there are several examples of successful transfer of PRRs to enhance disease resistance (Lacombe *et al.* 2010; Schoonbeek *et al.* 2015). PRRs or other components found in Arabidopsis might be transferred to crops or recombined with crop genes to link up with endogenous immune components and thereby engineer durable and broad-spectrum resistance (Ranf 2018). Indeed, MIK2 has many homologs in diverse crop plants, which may ease comparative genomics between model and crop towards interspecies functional complementation of *Fusarium* resistance.

## Material and Methods

### Plant material and growth conditions

*Arabidopsis thaliana* ecotype Columbia-0 expressing apoaequorin (Col-0^AEQ^) in the cytosol under control of the cauliflower mosaic virus 35S (CaMV35S) promoter was obtained from M. Knight (University of Durham, UK). The following mutant lines (Col-0 background) were obtained from the Nottingham Arabidopsis Stock Centre (NASC) and were genotyped using standard methods: *bak1-4* (TAIR accession code At4G33430; NASC accession code SALK_116202), *rbohD* (At5G47910; dSpm transposon mutant), *mik2-1* (At4g08850; SALK_061769), *mik2-2* (At4g08850; SALK_046987), *mik2-3* (At4g08850; GABI_208H02), *sobir1-12* (At2g31880; SALK_050715), *rfo1* (At1g79670; SALK_077975), *rfo2* (At1g17250; SALK_051677), *rfo3* (AT3G16030; SALK_040153), *rlp23-1* (At2g32680; SALK_034225) and *rlp42-1* (At3g25020; SALK_080324). *Cerk1-2* (*lyk1*; At3G21630; GK-096F09) was provided by Gary Stacey (University of Missouri, USA). *Mik2-like-1* (At1g35710; SALK_112341), *mik2-like-2* (At1g35710; GK-031G02), and double mutants *mik2-1/mik2-like-1* & *mik2-1/mik2-like-2* were provided by Cyril Zipfel (ETH, Zurich, Switzerland). The *rlp30-2* (At3g05360; SALK_008911) mutant line was provided by Thorsten Nürnberger (University of Tübingen, Germany); *herk1* (At3g46290; SALK_008043C), *the1-4* (At5g54380; SAIL_683_H03) and *wak2* (At1g21270; SAIL_12_D05) were provided by Kay Schneitz (TU Munich, Germany); *serk1-1* (At1g71830; SALK_044330), *serk2-1* (At1g34210; SALK_058020), *bkk1-1* (At2g13790, SALK_057955), *the1-1* (At5g54380; missense mutation G37D), *fer-4* (At3g51550; GABI_106A06) and *llg1-2* (At5g56170; SALK_086036) were provided by Martin Stegmann (TU Munich, Germany). The *bik1* (At2g39660; SALK_005291)*/pbl1* (At3g55450; SAIL_1236_D07) double mutant derives from Ranf *et al.* (2014). The line *mik2-3*^AEQ^ was generated by crossing *mik2-3* with Col-0^AEQ^ plants and was confirmed homozygous for both loci in the F_2_ generation using the appropriate primers. For growth in liquid medium, *A. thaliana* seeds were surface-sterilized then stratified for at least 48 hours (dark; 4°C) and grown in 24-well plates (15–20 seedlings per well) under long-day conditions (16 h of light; 20–22°C) in liquid MS medium (0.5x Murashige & Skoog medium plus vitamins (Duchefa, Haarlem, Holland)), 0.25% sucrose, 1 mM MES, pH 5.7). *A. thaliana* for ROS experiments were sown on a soil/vermiculite mix (8:1), stratified for 2 days (dark; 4°C) and grown under short-day conditions (8 h of light; 20°C– 22°C; 55% RH).

### *Fusarium* elicitor preparation, enrichment and characterization

*Fusarium graminearum* (isolate Fg006 from infected barley; Weihenstephan) is part of the TUM Chair of Phytopathology collection and *Fusarium oxysporum* (isolate DSM 62292) was provided by Ludwig Niessen (TU Munich, Germany). *Fusarium* isolates were grown in liquid malt medium (3% malt extract, 0.3% peptone) at 25°C on a 70 rpm shaker for 7-10 days. Fungal mycelium was collected using a fine mesh, thoroughly washed with distilled water, then lyophilized and ground using a pestle and mortar. For the preparation of FGE used for screening of EMS populations and comparable crude fractions from other *Fusarium* species, lyophilized mycelial powder was dissolved in water and autoclaved for 20 min at 121°C. The soluble fraction and up to two additional mycelial washes were pooled and lyophilized to concentrate elicitor activity then desalted at least twice using PD-10 desalting columns (GE Healthcare, Chicago, USA) until salt and other small molecules were removed. Crude elicitor fractions (i.e. FGE/FOE) were quantified based on the weight of starting material used (mg of lyophilized mycelium/mL). For the preparation of the enriched elicitor fraction (EnFoE), 10 g of lyophilized *F. oxysporum* mycelium was dissolved in 200 mL water and autoclaved as above. The soluble fraction was dialyzed in ⌀22 mm tubing with a 14 kDa cut-off (Biomol, Hamburg, Germany) overnight at 4°C and subsequently lyophilized. The powder was dissolved in 15 mL water, desalted to 30 mL then incubated with Macro-Prep High Q anion exchange media (Bio-Rad, Hercules, USA) in a 1:2 ratio (15 mL sample:30 mL beads) in 50 mL Falcon tubes on a shaker at RT for 1 h. The supernatant was collected and the beads were washed with 15 mL water for 10 min on a shaker in total 5 times. The resulting ‘Q_unbound_’ fractions were pooled and lyophilized; subsequently the powder was dissolved and desalted as above then incubated with Macro-Prep High S cation exchange media (Bio-Rad). The supernatant and five water washes were again collected and stored, but not used any further. The cation exchange medium was subsequently incubated 5 times with 15 mL 0.5 M NaCl on a shaker for 10 min, and supernatants of each elution step were collected, pooled and lyophilized (‘Q_unbound_/S_eluate’_). Finally, the ‘Q_unbound_/S_eluate_’ powder was dissolved in 15 mL water and desalted twice to a final volume of 60 mL EnFoE. Protein concentration was determined using Pierce™ BCA Protein Assay Kit (Thermo Scientific, Waltham, USA) according to the manufacturer’s instructions (stock concentration: 10 μg/mL). For proteinase treatment, elicitor samples were incubated overnight at 50°C with proteinase K (10 μg/mL) or without as a control and were heat-inactivated for 15 min at 95°C. To determine protein kinase dependency, Col-0^AEQ^ seedlings were pre-treated with K-252a (10 μM) or DMSO as a control within 96-well measurement plates 30 min before elicitor application.

### Elicitors

Chitin derived from shrimp shell (C9752; Sigma) was ground into a fine powder and resuspended in water for experiments (soluble fraction). The flg22 peptide (QRLSTGSRINSAKDDAAGLQIA), pep1 peptide (ATKVKAKQRGKEKVSSGRPGQHN) and Nep-1-like peptide from *F. oxysporum* (*Fo*NLP; AIMYAWYWPKDQPADGNLVSGHR) were synthesized on an Abimed EPS221 system (Abimed, Langenfeld, Germany).

### Detection of ROS in *A. thaliana* leaf discs

For detection of reactive oxygen species (ROS), ⌀4 mm leaf discs from 6-8 week old soil-grown plants were floated overnight on 200 μl water in 96-well plates (dark; RT). Shortly before the measurement, water was replaced with 100 μl of a 2 μg/ml horseradish peroxidase (type II; Roche, Rotkreuz, Switzerland) and 5 μM L-012 (WAKO Chemicals, Neuss, Germany) solution. Luminescence was recorded as relative light units (rlu) at 1-minute intervals with a Luminoskan Ascent 2.1 (Thermo Scientific) or a Tecan F200 (Tecan, Männedorf, Switzerland) luminometer. Measurements consisted of 10 min of background luminescence reading followed by a further 60 min after elicitors were added to the appropriate final concentration. Results were normalized to average ROS at 5 min before treatment followed by subtraction of values for untreated controls (included for each genotype on the same plate). Individual measurements over the 60 minutes (kinetics) or the highest luminescence reading achieved during this time (maxima) were used for analyses. To account for plate effects and standardize replicate measurements, values were normalized to the mean rlu measured in WT plants (set to 1) on respective plates (indicated as: norm. ROS).

### Aequorin luminescence measurements

For measurement of aequorin luminescence, 8-10 day old liquid-grown apoaequorin-expressing seedlings were placed individually in 96-well plates containing 100 μl of 5 µM coelenterazine-h (PJK, Kleinblittersdorf, Germany) in water overnight (dark; RT). Resting luminescence levels were determined by scanning each well 12 times in 10 s intervals, then a 25 µL elicitor preparation was added at the appropriate concentrations and luminescence was measured 180 times in 10 s intervals (Luminoskan Ascent 2.1). The remaining aequorin was discharged by the addition of 150 μl discharge solution (2 M CaCl_2_, 20% ethanol) per well and measured as L_max_. [Ca^2+^]_cyt_ concentrations were calculated as L/L_max_ (luminescence counts per second relative to total luminescence counts remaining) as described (Ranf, *et al.*, 2012). For [Ca^2+^]_cyt_ measurements in different plant tissues, 9 day-old seedlings grown vertically on solid MS medium (as described above with addition of 0.9% agargel; Sigma, St. Louise, USA) were either dissected into roots and above ground tissues with a scalpel or left intact and respective tissues were incubated in substrate as indicated above. Individual measurements over the 30 minutes (kinetics) or the highest luminescence reading achieved during this time (maxima) were used for analyses. Normalization to WT plants was carried out in the same way as described for ROS measurements (indicated as: norm. [Ca^2+^]_cyt_[L/L_max_]).

### Immunoblot analysis of MAPK phosporylation

Analyses of phosphorylated MAPKs were performed as previously described (Ranf *et al.* 2011). Briefly, 14 day old liquid-grown seedlings were equilibrated for 24 h in fresh MS medium. The elicitor solutions were later added at the appropriate concentrations and seedlings were harvested at the stated time points. Protein extraction and immunoblotting with an antibody against phosphorylated MAPKs (p44/42; 9101; Cell Signaling Technology) was then performed. Blots were visualized using SuperSignal West Dura Extended Duration Substrate (Thermo Scientific) in a Fusion SL Imager (Vilber Lourmat, Eberhardzell, Germany) with the accompanying software. Amido Black staining was performed to confirm equal protein loading and blotting (Rubisco band of approximately 56 kD is shown).

### Gene-expression analysis

10 day old liquid-grown seedlings were equilibrated in fresh MS medium for 24 h before water or elicitor treatment at the stated final concentrations. Whole seedlings were harvested at 2 h and total RNA was extracted using the conventional Trizol/chloroform method, followed by treatment with DNase I (Thermo Scientific) and then reverse-transcription using oligo(dT) and RevertAid reverse transcriptase according to the manufacturer’s instructions (Thermo Scientific). Complementary DNA (from 1-2 µg RNA) was diluted 1:10 in water and 3 μl per reaction was mixed together with SYBR Green™ (Thermo Scientific) and gene-specific primers (Supplementary Table 1) to a final volume of 20 μl in clear 96-well plates with clear plastic lids (Agilent Technologies, Waldbronn, Germany). QRT-PCR assays were performed using an AriaMx Real-Time PCR system (G8830A instrument; Agilent) and the accompanying software (version 1.5). Amplification conditions were as follows: 40 cycles of 30 s at 95°C, 30 s at 60°C and 30 s at 72°C. The SYBR-specific fluorophore was quantified during the reaction by the instrument and threshold cycles values calculated according to the software instructions. Data were exported and analyzed in MS Excel 2010. The expression of target genes was calculated using the 2^−ΔΔCT^ method as previously described by Livak and Schmittgen (2001) and normalized to expression of the house-keeping gene *ubiquitin*. The mean values from two technical replicates included for each cDNA–primer combination were used for analysis. Data represent the relative expression of each target gene compared to the water-treated control for each genotype.

### *Arabidopsis thaliana* mutant screen

The Col-0^AEQ^ EMS mutant screening procedure was similar to previously described (Ranf *et al.* 2012; Ranf *et al.* 2015) with some minor modifications. Col-0^AEQ^ seeds underwent mutagenesis for 18 h at room temperature with gentle shaking in 0.3% ethyl methane sulfonate (EMS; Sigma). Treated seeds were washed with a 5% sodium thiosulfate solution to inactivate EMS, rinsed several times with water and dried. M_1_ plants were grown on soil and seeds were harvested from individual plants. Approximately 12-24 M_2_ seedlings per M_1_ plant were measured for aequorin luminescence in response to FGE as described above but without discharge. Seedlings with a markedly lower or later elevation in aequorin luminescence than that of control Col-0^AEQ^ seedlings were ‘rescued’ from the 96-well plates to solid MS media for approximately one week then transferred to soil for seed setting. Mutant phenotypes were verified by analysis of the corresponding M_3_ offspring with quantitative [Ca^2+^]_cyt_ measurements. The *fere1* mutant line was backcrossed (Col-0^AEQ^) in order to reduce background mutation levels and phenotypic analysis of F_1_ and F_2_ populations resulting from this cross demonstrated a single recessive locus. A single F_2_ offspring homozygous for *fere1* was used for all experiments and subsequent genetic material produced in this study.

### Mapping and sequencing of candidate genes

Mapping populations were generated by crossing *fere1* with the *A. thaliana* ecotype Landsberg *erecta*-0 (L*er*-0). Approximately 200 F_2_ offspring were grown on soil for seed setting. As outcross populations introduce ecotype-associated phenotypic variation, which makes accurate assessments of mutant phenotypes more difficult, F_3_ progeny (approx. 24 per F_2_) were phenotyped using quantitative [Ca^2+^]_cyt_ measurements in order to better determine *fere1* homozygosity. Additionally, the EnFOE fraction was used in order to better distinguish between WT-like and *fere1*-like phenotypes. F_3_ plants were grown out on soil and harvested at 2-weeks for genomic DNA isolation in order to reconstitute the genetic make-up of each F_2_ parent. Finally, 22 *fere1*-like F_2_ and 5 WT-like F_2_ individuals were used for mapping. The analysis of PCR-based markers localized *FERE1* to a mapping interval of approximately 0.4 MB (position. 5372922-5756738) on chromosome 4 containing 121 gene loci (TAIR; https://www.arabidopsis.org/). Finer mapping was not necessary as the *MIK2* gene (position. 5636489-5640636) is the only gene within this interval that encodes a candidate component of a PRR complex by protein domain architecture. For sequence analysis, full-length *MIK2* was amplified from genomic DNA by PCR with gene-specific primers using Phusion Hot Start High-Fidelity DNA Polymerase (Thermo Scientific) and purified PCR products were sequenced using appropriate nested primers. A SNP resulting in the W876STOP mutation in *fere1* was confirmed and a cleaved amplified polymorphic sequence (CAPS) marker was developed to genotype for the mutation. For genotyping by CAPS, genomic DNA samples were amplified with the appropriate primers (Supplementary Table 1) using BioTherm™ DNA polymerase (AxisBio Science Genecraft, Köln, Germany) followed by a 2 hour digestion at 37°C with the enzyme HpyF3I (DdeI) (Thermo Scientific) according to the manufacturer’s instructions. Digestion products were visualized on a 2% TAE gel. For reciprocal crossing experiments, the independently obtained T-DNA insertion line of *mik2-3* (see above) was found to be allelic to *fere1* in a complementation test; i.e., F_1_ hybrids resulting from reciprocal crosses failed to complement each other for EnFOE-induced ROS production.

### Molecular cloning and generation of transgenic lines

To generate a MIK2 construct with a c-terminal cMyc fusion (EQKLISEEDL) under the control of its own promotor, the full-length genomic fragment of *MIK2* (At4g08850) without the stop codon and including a 2 KB upstream region containing the putative promotor was first amplified from Col-0^AEQ^ genomic DNA using appropriate primers (Supplementary Table 1). The pENTR™/D-TOPO™ Cloning Kit was used (Thermo Scientific) to generate a Gateway compatible entry clone then the gene was reshuffled into the pEarleyGate 303 destination vector *via* a standard LR cloning method. To generate a construct containing the native MIK2.1 splice form, a GoldenGate cloning strategy was used. The MIK2 promotor sequence described above and MIK2 cDNA sequence were sub-cloned into respective modules using appropriate primers (Supplementary Table 1) and assembled together with the CaMV35S terminator sequence into an appropriate binary vector for plant expression. Consequently, constructs were transferred into *Agrobacterium tumefaciens* strain GV3101 and *fere1* plants were transformed by floral-dip transformation. Transgenic plants were selected by BASTA spraying on soil (glufosinate-ammonium; Bayer CropScience, Langenfeld, Germany) and independent homozygous T_3_ lines were later established by BASTA selection on MS agar plates.

## Funding

This work supported by the BASF Plant Science Company GmbH (research and development cooperation with S.R. and R.H.) and a grant from the German Research foundation to R.H. (HU886/11).

## Competing interests

The authors declare no competing interests.

## Contribution of the authors

R.H. and S.R. designed the study. S.R., R.H., and A.D.C. planned the experiments and interpreted results. A.C., L.R. and J.M. optimized methodology and performed experiments. A.D.C. and J.M. prepared figures. A.D.C. and R.H. wrote and revised the manuscript.

## Acknowledgements

We are grateful to Dr. Holger Schultheiss (BASF Plant Science Company GmbH) for fruitful discussions and to Dr. Martin Stegmann (Phytopathology, TU Munich) for critical reading of the manuscript.

---

**Figure S1:**
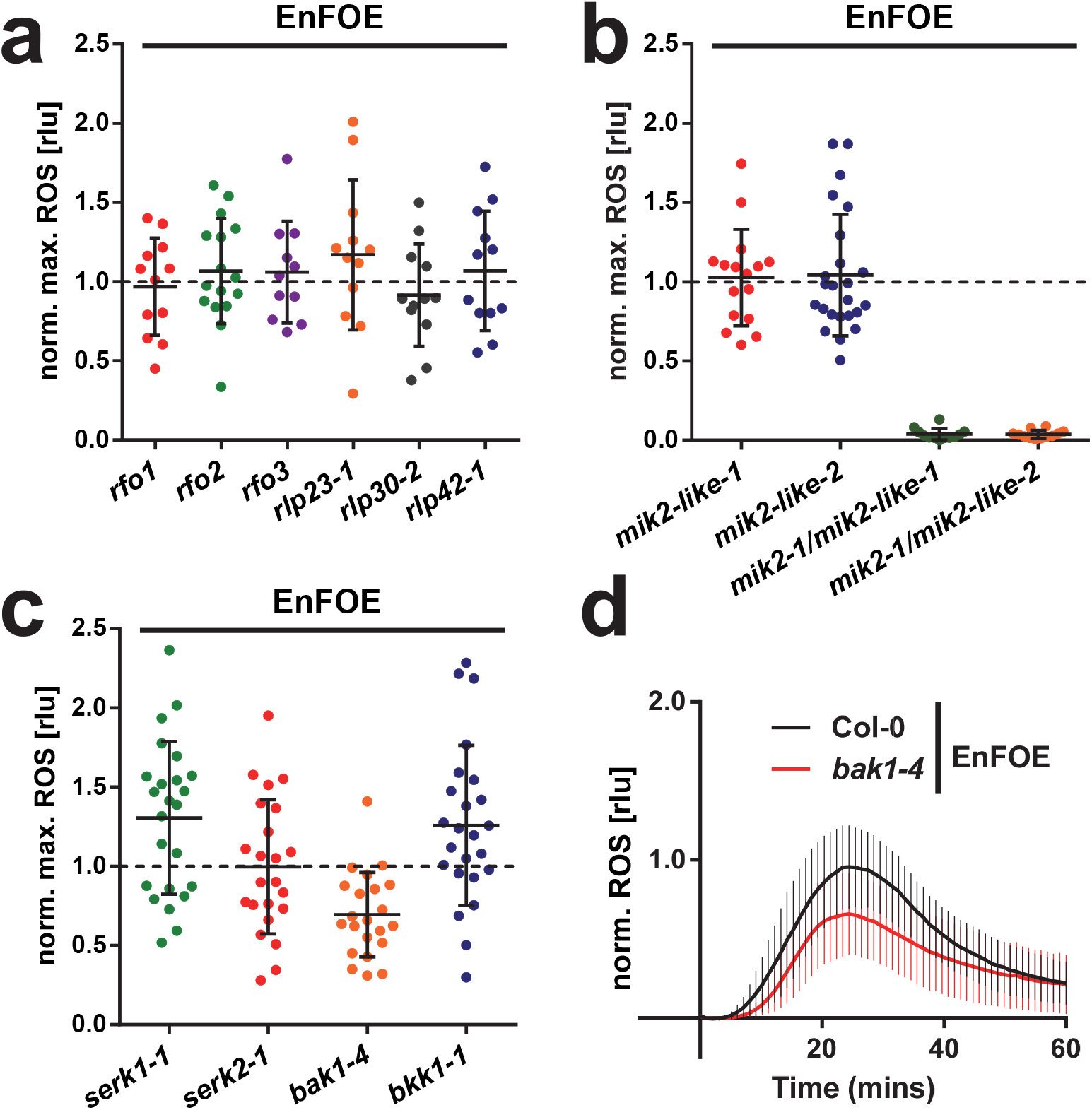
Additional ROS response measurements in relevant Arabidopsis genotypes. [**a-c**] Maximum ROS accumulation measured in individual leaf discs (each plotted point) of indicated mutant genotypes over 60 minutes after elicitation with EnFOE (0.5 µg/mL) normalized to the mean maxima of Col-0 plants (set at 1; dotted line) with error bars representing s.d. (*n*=12–24). [**d**] ROS accumulation kinetics in Col-0 and *bak1-4* leaf discs after elicitation with EnFOE (0.5 µg/mL). Data represent mean relative light units [rlu] measured from individual leaf discs normalized to respective values from Col-0 plants with error bars representing s.d. at each time point (*n*=22–23). Data shown (a-d) is pooled from at least three independent biological replicates.

**Figure S2:**
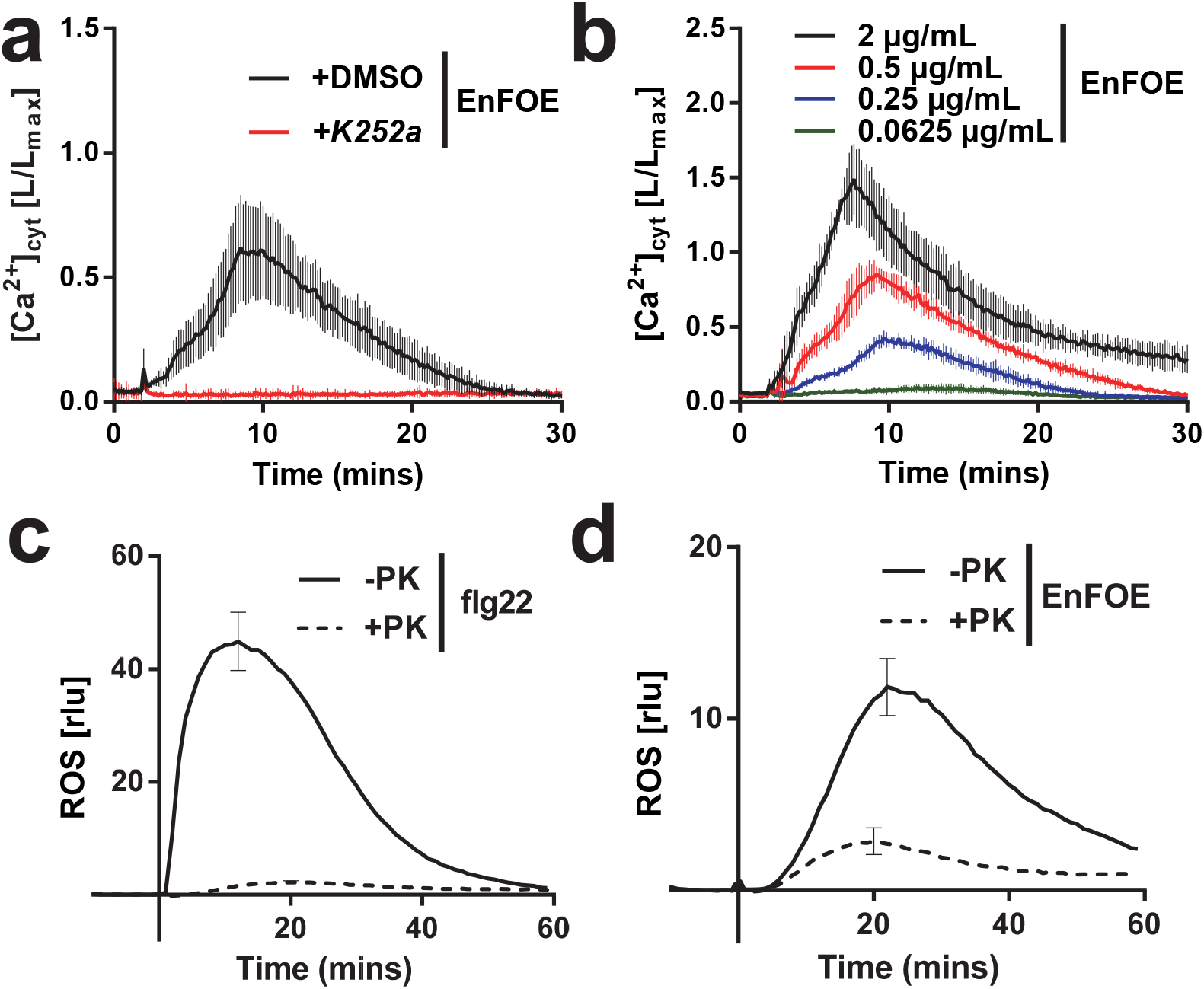
Characterisation of *Fusarium* elicitor responses. [**a**] Cytosolic Calcium quantification ([Ca^2+^]_cyt_) kinetics in Col-0^AEQ^ seedlings after elicitation with the enriched *F. oxysporum* elicitor fraction (EnFOE; 0.5 µg/mL) with or without pre-treatment with the kinase inhibitor K252a. Data represent mean values measured in individual seedlings with error bars representing s.d. at each time point (*n*=12).[**b**] Cytosolic Calcium quantification ([Ca^2+^]_cyt_) kinetics in Col-0^AEQ^ seedlings after elicitation with indicated concentrations of EnFOE. Data represent mean values measured in whole seedlings with error bars representing s.d. at each time point (*n*=4). [**c-d**] ROS accumulation kinetics in Col-0 leaf discs after elicitation with [c] flg22 (200 nM) or [d] EnFOE (0.5 µg/mL) with or without elicitor pre-treatment with proteinase K (PK). Data represent mean relative light units [rlu] measured from individual leaf discs with error bars representing s.d. at the maximum of the curve (*n*=4). Data shown is representative from at least three independent experiments with similar results (b-d) or normalized pooled data from three independent biological replicates (a).

**Figure S3:**
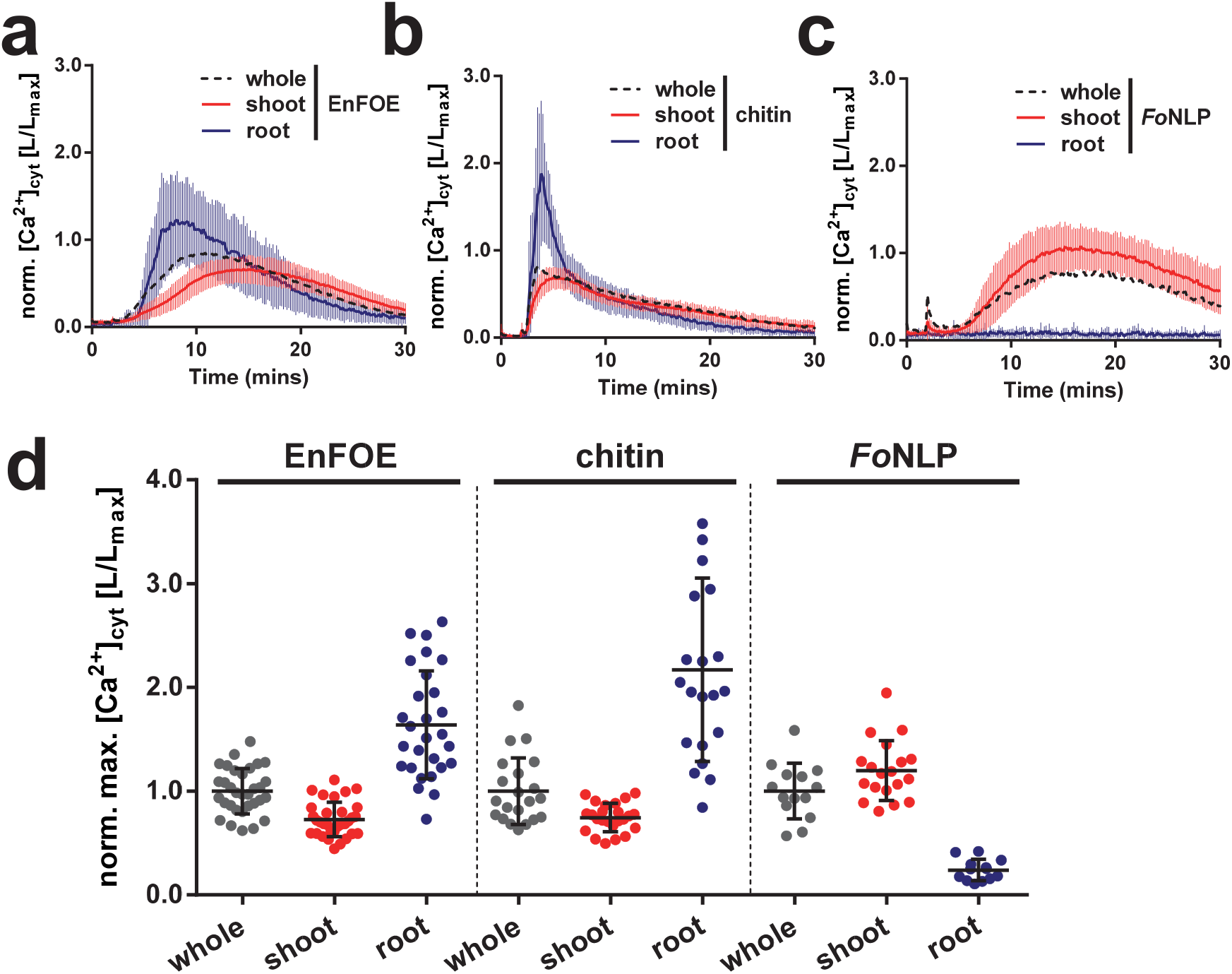
Root versus shoot responses to fungal elicitors. **[a-c**] Cytosolic Calcium quantification ([Ca^2+^]_cyt_) kinetics in whole Col-0^AEQ^ seedlings, or dissected shoot/root tissues after elicitation with [a] EnFOE (0.5 µg/mL), [b] chitin (75 µg/mL) or [c] *Fo*NLP (1 µM). Data represent mean values measured in indicated tissues normalized to respective values from whole Col-0^AEQ^ seedlings (dotted line) with error bars representing s.d. at each time point (*n*=12–32). [**d**] Maximum [Ca^2+^]_cyt_ measured in whole Col-0^AEQ^ seedlings or dissected shoot/root tissues (each plotted point) over 30 minutes after elicitation with EnFOE (0.5 µg/mL), chitin (75 µg/mL) or *Fo*NLP (1 µM) normalized to the mean maxima of whole Col-0^AEQ^ seedlings (set at 1) with error bars representing s.d. (*n*=12–32). Data shown (a-d) is pooled from at least three independent biological replicates.

**Figure S4:**
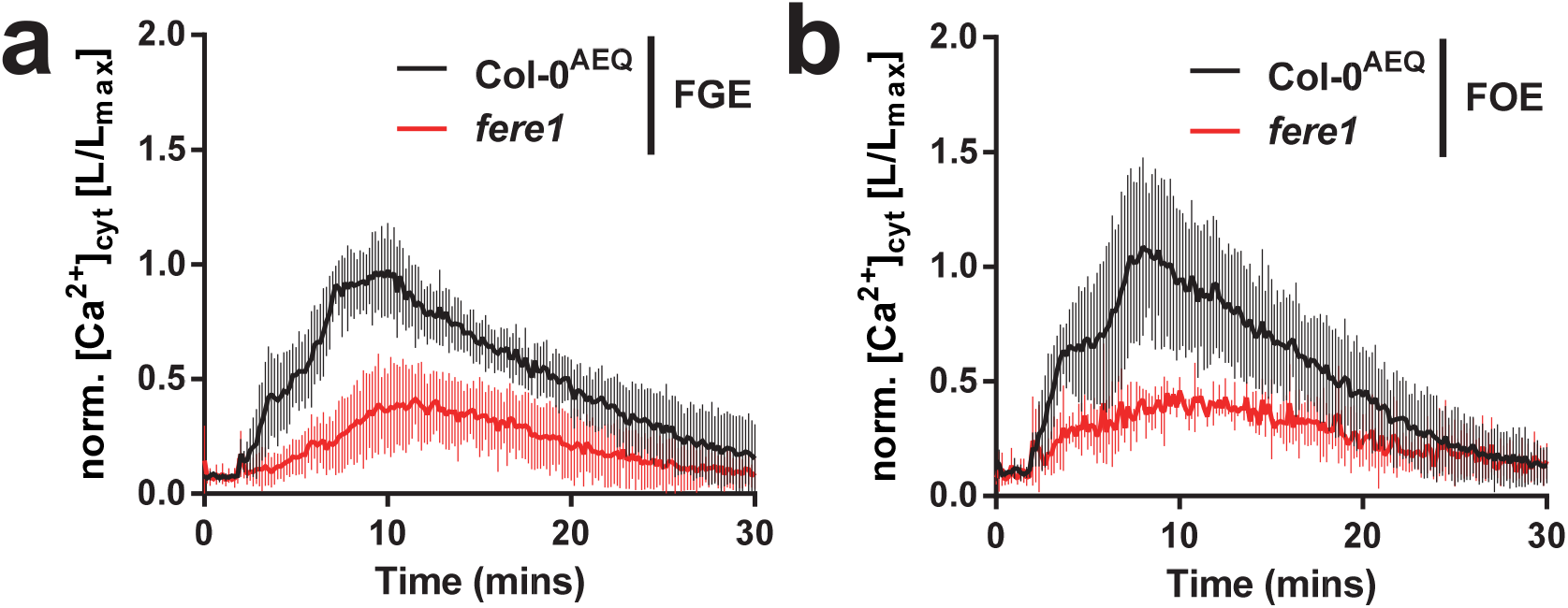
*Fusarium* crude elicitor responses in wild type and *fere1* mutants. [**a-b**] Cytosolic Calcium quantification ([Ca^2+^]_cyt_) kinetics in Col-0^AEQ^ or *fere1* seedlings after elicitation with [a] crude *F. graminearum* elicitor or [b] *F. oxysporum* elicitor fractions (FGE/FOE; derived from 1 mg mycelium/mL). Data represent mean values measured in indicated genotypes normalized to respective values from Col-0^AEQ^ seedlings with error bars representing s.d. at each time point (*n*=4). Data shown (a-b) is representative from at least three independent experiments with similar results.

**Figure S5:**
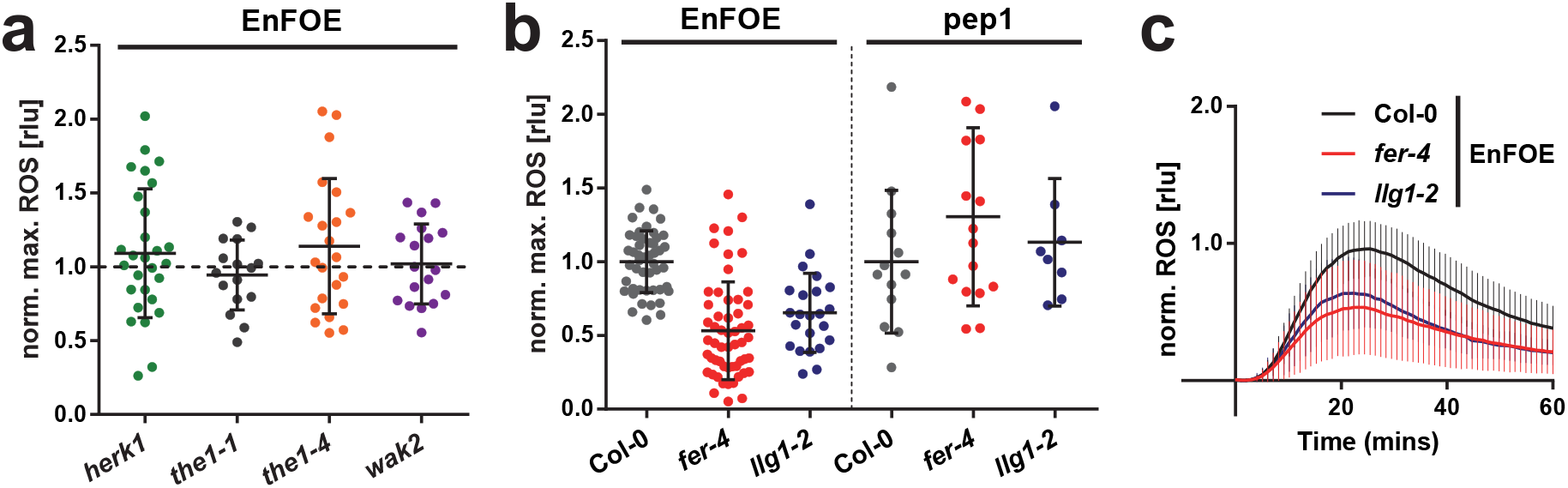
Elicitor responses of cell-wall integrity and related mutants. [**a-b**] Maximum ROS accumulation measured in individual leaf discs (each plotted point) from indicated mutant alleles after elicitation with the enriched *F. oxysporum* elicitor fraction (EnFOE; 0.5 µg/mL) or Arabidopsis pep1 (1 µM) normalized to the mean maxima of Col-0 plants (set at 1; dotted line in a) with error bars representing s.d. (*n*=8–55). [**c**] ROS accumulation kinetics in wild type Col-0 and indicated mutant leaf discs after elicitation with EnFOE (0.5 µg/mL). Data represent mean relative light units [rlu] measured from individual leaf discs normalized to respective values from Col-0 plants with error bars representing s.d. (*n*=23–55). Data shown (a-c) is pooled from at least two independent biological replicates.

**Supplementary table 1:**
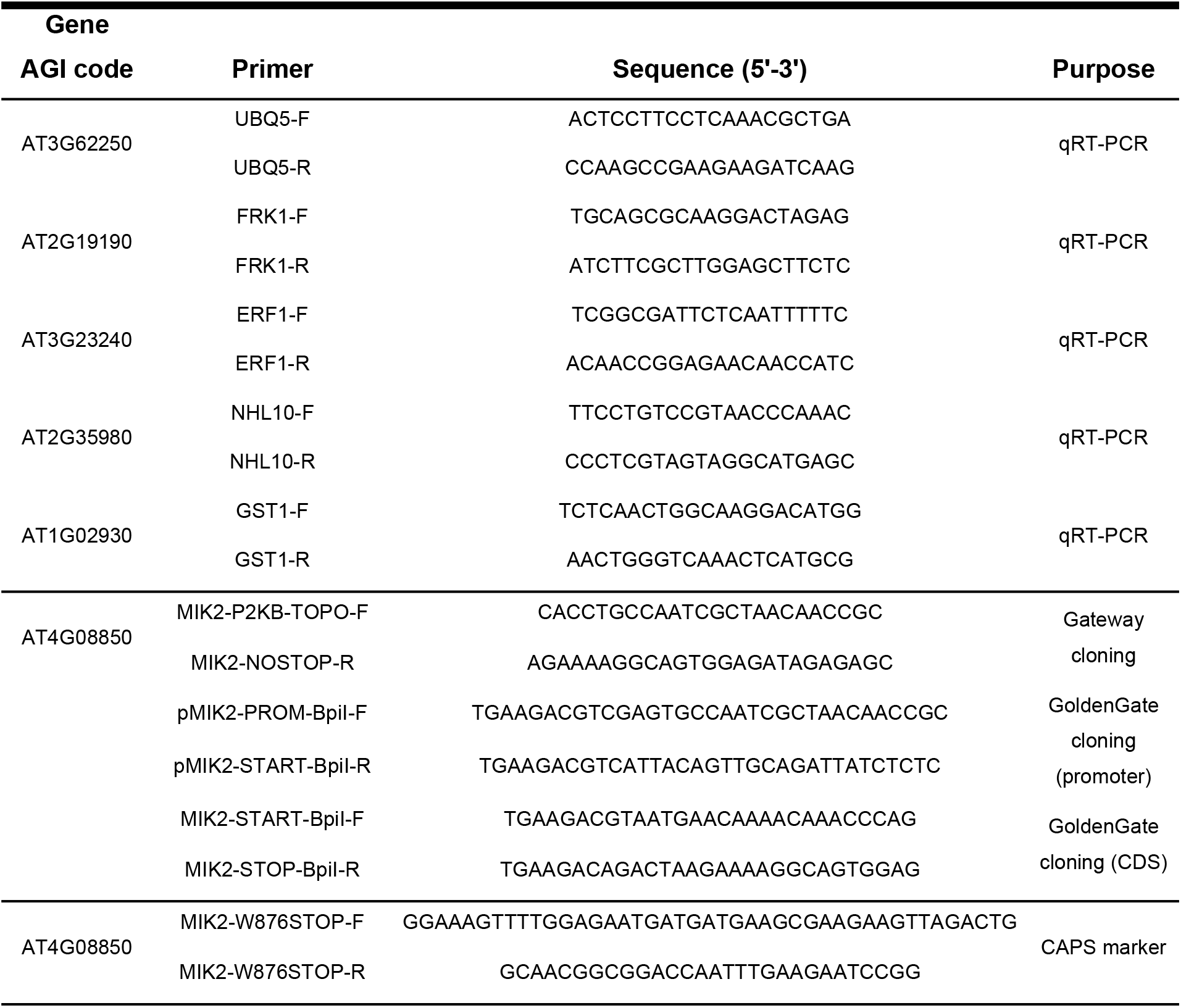
Primers used in this study.

